# The impact of fibronection stripe patterns on the cellular and nuclear morphology of fibroblasts

**DOI:** 10.1101/302687

**Authors:** Pooya Mamaghani, Athene M. Donald

**Affiliations:** Department of Physics, University of Cambridge, Cavendish Laboratory, 19 JJ Thomson Avenue, Cambridge CB3 0HE, UK

**Keywords:** Cell adhesion, Substrate texture, Cellular volume, Cell thickness, Nuclear deformation, Perinnuclear actin cap

## Abstract

The effect of biochemical environmental signals on cell mechanisms has been the subject of numerous studies for a long time. However, the *in-vitro* studies of biophysical cues on cells and tissues have recently become a popular focus of research. The development of micro-fabrication techniques has allowed the study of certain aspects of cell-substrate interactions in a more detailed form. Micro-topographical patterns on the cell substrates have been used to study many cell functions such as cell migration, adhesion, gene expression, cell division and differentiation. An understanding of cell-substrate interactions and the potential ability to control the interactions have very important applications in the field of tissue engineering and regenerative medicine. We have fabricated ridge-groove micro patterns on polydimethylsiloxane (PDMS) substrates with different ridge widths (8μm, 10μm, 12 μm, 25μm and 50μm) using standard photolithography technique. We used these patterns to print fibronectin stripes on PDMS substrates. NIH/3T3 fibroblast cells were cultured on these stripes and the dynamics of morphological changes were monitored in steady spreading phase (S-phase). Our data revealed that the thickness of the cell, measured by confocal microscopy, is considerably larger (approximately 40%) among the cells spreading on narrower stripes (8μm, 10μm and 12μm) compared to the cells expanding on wider (including control) patterns. The number of perinuclear actin stress fibers is significantly lower among narrower stripes which probably explains the cell thickness results. Confocal microscopy revealed that the cellular volume increases during cell adhesion processes and volume increase is positively correlated with the width of stripes. Nuclear volume also increases considerably during cell adhesion; however, confining cells on fibronectin stripes reduces nuclear volume enlargement independent from the of stripe size.

## Introduction

Some cell lines are capable to adhere to the surface (substrate), extracellular matrix (ECM) or the adjacent cells with the facilitation of a complex set of cell adhesion molecules (CAMs). Cell adhesion plays a critical role in structural integrity of tissues, maintaining multicellular structures, pathogenesis of infectious organisms and signal transduction (1). Adherent cells, such as fibroblasts, demonstrate the ability to conform to the external topographical cues and move in accordance to the texture of substrate. This phenomenon was first observed in 1912 (Harrison, 1912) and later was coined “contact guidance’’(2). Contact guidance occurs not only *in vitro* when the cells are in contact with a two dimensional substrate texture, it also frequently happens in the native environment of the cells where they are constantly exposed to topographical cues in the form of the extracellular matrix (ECM) and surrounding cells ((3);(4)). As a result, studying the phenomenon is of prominent importance in understanding several key processes such as wound healing, embryogenesis, nerve regeneration, angiogenesis and stem cell differentiation ((5);(6);(7);(8);(9)). 2D topographic patterns printed by adhesive molecules such as fibronectin, are optimized models with precisely controlled parameters to mimic cell-ECM and cell-cell interaction in the native environment of cells. These *in vitro* models are increasingly capturing the attention of researchers, as accumulating evidence reaffirms that physical interaction of the cell and extracellular environment (either in the form of topography or matrix stiffness, or the combination of both) plays a critical role in modulating vital cell machineries such as cell proliferation, gene expression, differentiation and signal transduction and mechanosensing ((10);(11);(12);(13)(14);(15)).

Our fibronectin stripes were printed from ridge-groove patterns. Ridge-groove arrays are a family of topological patterns that have been frequently used for studying contact guidance since 1979. The patterns were first introduced to explain previously observed contact guidance among chick heart fibroblast and kidney epithelium cells induced by fine grooves in plastic culture dishes(16). The patterns mimic some of frequently occurring situations in the living microenvironment of the adherent cells such aligned collagen fibres. The majority of the cells lines that have been cultured on these patterns tend to align themselves along the major axis of the grooves (17). Depending on the dimensions of the ridges and grooves, adherent cells show characteristic behaviour such as bridging over the grooves without contacting their surface (bridging), cell confinement, cell traversing (connecting) between ridge and grooves ((16);(18)). Micro stripes patterns printed by CAMs such as fibronectin or collagen using ridge-groove patterns as the template have frequently been employed for quantitative study of the dynamics of cell migration and adhesion, force coupling between the nucleus and adhesion complexes of the cytoskeleton, cytoskeletal filaments, adhesion morphology and mesenchymal stem cell differentiation ((19);(20);(21);(22);(23)).

Although 2D morphological features of adherent cells on these patterns has been well studied, due to the more complex nature of 3D microscopy, the 3D behaviour of these cells are poorly understood. It is known that when fibroblast cells are seeded on fibronectin-printed stripes, the cell elongates itself along the major axis of the pattern(19). Usually the nucleus of the elongated cells also get elongated (24) due to the compressive pressure applied by cytoskeletal microtubule motors and the compressive force applied by actin stress fibers(25). However, the extent of nucleus elongation can exhibit multimodal behaviour among some cell lines such as human mesenchymal stem cells (hMSCs), so that the histogram of aspect ratio of the hMSCs shows two distinct groups of elongated and round nucleus. (26). Because of the pivotal role that nuclear deformation plays in the behaviour and fate of cells, substantial interest has developed exploring this opportunity to control the deformation using substrate topography((27);(28);(29)), particularly for control of proliferation and differentiation of stem cells((28);(10);(30)). More recent studies have even shown that certain topographical cues on the substrate can effectively be used to optimize reprograming of fibroblasts to induce neurons(31).

It has been shown that the projected area of fibroblasts adhering to the narrower fibronectin stripes, decreases significantly(19). Furthermore, total cellular volume and nuclear volume of fibroblasts does shrink when the cells fully spread on the substrate(25). However, a precise quantified study of the effect of adhesion on the cellular thickness and volume among the fibronectin stripe pattern has not yet been performed. This is particularly important as we know that the geometry of cytoskeletal stress fibers plays a key role in deformation and bifurcation of the cell nucleus and this directly links to the gene regulatory machinery of the cell((25);(32);(27)). As a result, it is crucial to acquire a clear 3D image of both cytoskeletal and nuclear deformation during the adhesion process in a model system as it will help us to gain better understanding of possible gene regulatory effects of extracellular microenvironment.

## Materials and Methods

### Cell culture

NIH/3T3 mouse embryonic fibroblasts (ATCC, Middlesex, UK) were cultured in a medium consisting of Dulbecco’s modified Eagle’s medium (DMEM) (Invitrogen, Paisley, UK) enriched with 10%(v/v) fetal bovine serum (Sigma-Aldrich, Dorset, UK), 100 μg ml^−1^ of streptomycin and 100 units/ml of penicillin and (Sigma-Aldrich), at 37 °C and 5% CO2.

### Soft Lithography

The template pattern to generate PDMS stamps for micro-contact printing was fabricated using a photolithography technique (33). First, the photoresist SU-8 2015 liquid (Chestech Ltd, Rugby, UK) was coated on a single side polished (SSP) type P silicon wafers (thickness: 320-350 μm, diameter: 50.8 mm, resistance: 0-100 ohm-cm) (University Wafers, Massachusetts, USA).Afterwards, SU-8 coated silicon wafers were baked on the hot plate at 60 °C for 1 minute and at 95 °C for 2 minutes subsequently. Wafers were left to cool down and placed in a mask aligner (Karl Suss MJB4, Garching, Germany) and put into soft contact with the chromium mask that contains desired strip pattern (Compugraphics, Glenrothes, UK) afterwards. Wafers were exposed to UV light for 4 seconds and baked on the hot plate for 1 minute at 60 °C and for 3 minutes at 95°C subsequently in order to crosslink the exposed SU-8. Cooled wafers were immersed in SU-8 developer (Chestech Ltd, Rugby, UK) for 1 minute in order to remove un-cross linked SU-8 polymers. Prepared template were rinsed twice with isopropyl alcohol, dried with pressurized nitrogen and baked at 150°C for 10 min. A layer with approximately 5 mm of PDMS1 was poured onto prepared SU-8 patterns. Wafers were put into vacuum chamber to remove trapped bubbles and baked at 150°C for about 15 minutes to cure PDMS.

### Substrate preparation

Substrate preparation and micro-printing works carried out mostly in accordance with the conventional method (34) with some adjustments for our system. PDMS was poured onto a circular coverslip (diameter: 22 mm, thickness: 0.19mm). The coverslip was pre-rinsed with isopropanol, sonicated for 10 minutes in an ionized water bath and blow-dried with pressurized air in order to clean it. Afterwards, the coverslip was spin coated at 1700 rpm for 35 seconds in order to fabricate a 55μm layer of PDMS. This PDMS coated coverslip was baked at 150°C for 15 minutes for curing. A narrow circular line of uncured PDMS was streaked on the edge of a custom-made metal ring and the pre-prepared coverslip was mounted on the ring. The coverslip mounted ring was baked at 150°C for 15 minutes to cure the PDMS glue.

### Micro-contact Printing

The desired PDMS stamp was cut from the patterned PDMS film. The cut PDMS stamp was sterilized by immersion in 70% ethanol. Afterwards, the ethanol was washed away gently from the stamp with 1x autoclaved PBS. PDMS stamps were exposed to 50μg/mL fibronectin from bovine plasma (Sigma-Aldrich) and were incubated at 37°C for 1 hour. Stamps were gently rinsed three times with autoclaved 1x PBS and once with autoclaved ionized water subsequently. The PDMS substrate embedded in the custom-made metal ring was treated with oxygen plasma treatment inside a plasma cleaner machine (FEMTO plasma cleaner, Diener Electronic, Ebhausen, Germany) at 90W and 25 Standard cubic centimetres per minute (Sccm) flow rate for 20 seconds before being stamped with a fibronectin-inked PDMS stamp. Oxygen plasma treatment enhances pattern transfer in the stamping process by increasing the hydrophilicity of the PDMS substrate (35). The PDMS stamp was put in contact with the PDMS substrate and a gentle pressure, using a standard mass (~50-100g) was applied in order to transfer the pattern to PDMS substrate. The substrate-stamp complex was incubated at 37°C for 3 hours subsequently. Afterwards, the stamp was removed and the PDMS substrate was rinsed twice with 1x PBS and twice with ionized water. The PDMS substrate was moved and kept in 4°C fridge in the end of the process.

### Cell culturing on the patterns

Cells were allowed about 200 minutes to spread on fibronectin-coated patterns and confocal microscopy was performed in the z direction (Fig. 1). Previous temporal studies on the various adherent cells have shown that the projected cell area increases over time exhibiting a three phase sigmoid curve((36),(37),(38),(39),(40)) which is very similar to the same temporal dynamic of cytoskeletal filamentous actin polymerization(41), the critical mechanism behind cell spreading (38). Area measurements of 3T3 fibroblast cells have illustrated that by 200 minutes after seeding on fibronectin coated PDMS substrates the cell area reaches a steady state phase(S-phase) after an initial lag slow growth (L-phase) followed by a rapid area expansion phase (E-phase)(19).

**Fig. 1:**
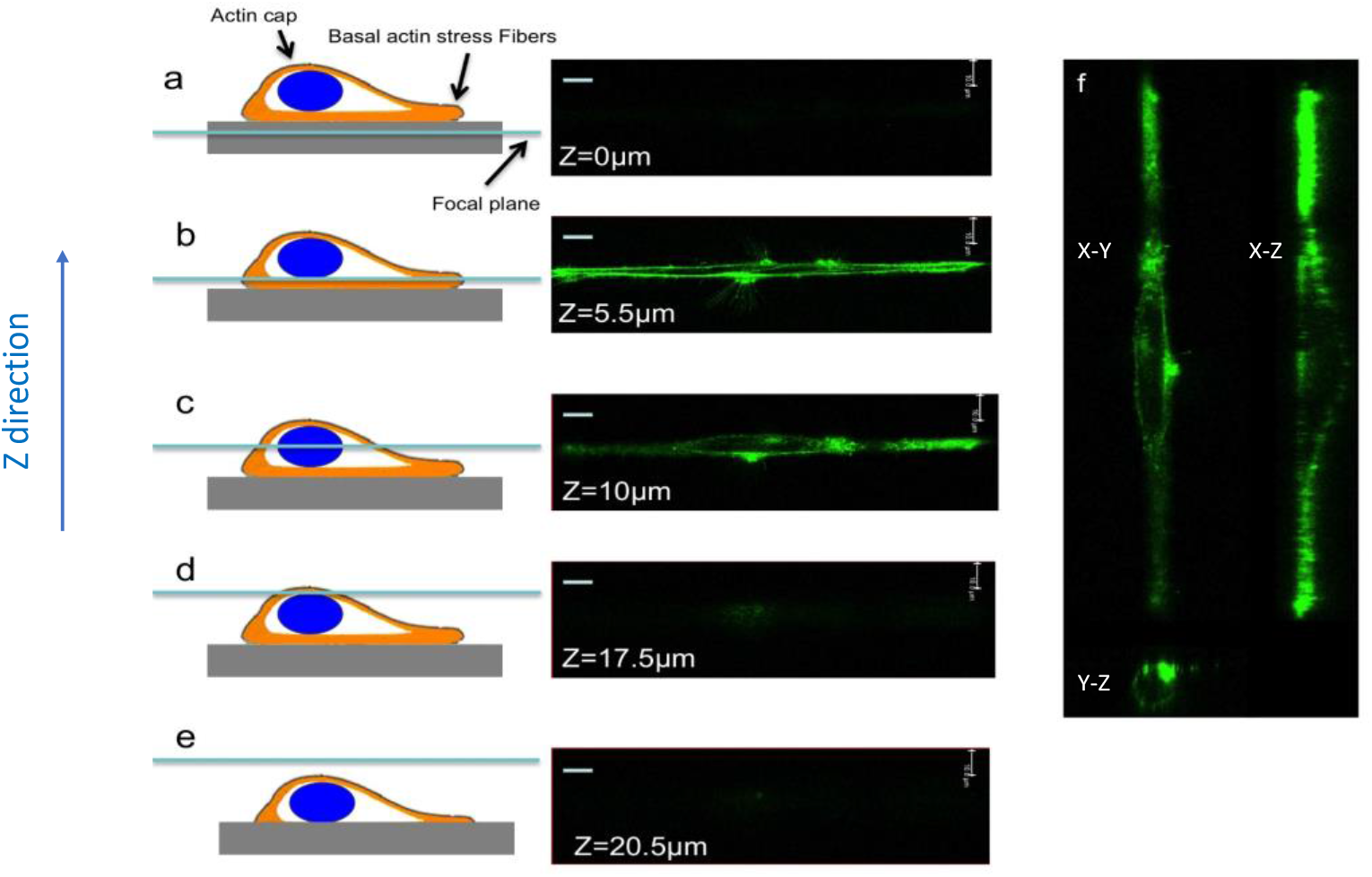
Confocal microscopy of cells on fibronectin stripes. Focal plane inside substrate(**a**), basal stress fibers(**b**) across the nucleus(**c**), perinuclear actin cap(**d**) and outside the cell(**e**). Scale bars 10μm.

### Fluorescence microscopy

#### Filamentous actin microscopy

16% formaldehyde stock (Polysciences Inc., Eppelheim, Germany) was diluted to 4% with 1xPBS. Cells were fixed with 4% formaldehyde for 10 minutes at room temperature and gently rinsed with PBS twice. Afterwards, the cells were soaked in 1x PBS with 0.1 % Triton X-100 (Sigma-Aldrich, Dorset, UK) for 3 minutes. The cells were again rinsed twice with 1x PBS and exposed to 1% BSA blocking solution (Sigma-Aldrich, Dorset, UK) for 30 minutes. The cells were stained with 6.6 μM Alexa Fluor 488 Phalloidin (Life Technologies, Paisley, UK), incubated for 30 minutes at room temperature and rinsed twice with 1x PBS afterwards. Stained cells were covered with aluminium foil and kept at 4°C. An inverted confocal scanning laser microscope (Leica TCS SP5, Wetzler, Germany) was used for fluorescent microscopy. A 100X oil immersion objective lens with 1.4 Numerical Aperture were typically used. Type F 1.518 oil from Leica was used as the immersion oil. Images were analysed with the LAS AF Lite software suite from Leica. For volume estimation of detached cells, cells were treated for 3 min at 37°C with Trypsin-EDTA (0.25% Trypsin, and 0.53 mM EDTA) (Sigma-Aldrich) and then re-suspended in fresh medium and were quickly imaged were 40X bright-field microscope. Volumes were estimated with the assumption of spherical geometry for suspended cells.

#### Cellular Thickness measurement

The high precision stage control in z direction (10nm), leaves the diffraction-limit as the only resolution limiting factor. Thisis about 0.2 μm for an objective numerical aperture (NA) of 1.4 and 488nm excitation light. With such a resolution we were easily able to visualize basal actin stress fibres (Fig. 1(b)), nucleus region(Fig. 1(c)) and pronuclear actin cap(Fig. 1(d)).

#### Nuclear microscopy

Cells were fixed with 4% formaldehyde and gently rinsed with 1xPBS three times. Cells were covered with 1μM of TO-PRO^®^-3(Life technologies, Paisley, UK) solution in 1xPBS and incubated for 30 minutes at room temperature protected from light. Subsequently, the staining solution was removed and cells were rinsed three times with 1xPBS. Fluorescent microscopy was performed using the same setup as filamentous actin with 632nm He-Ne excitation laser and emitted light was captured around 661nm. For the measurement of nuclear volume of suspended cells, the medium was removed after trypzination and cells were re-suspended in 1% PBS. Cells were stained with 5μg/mL Hoechst^®^ 33342 (Thermofisher scientific, Paisley, UK) for 10 min and quickly were imaged. Spherical assumption were considered to estimate to nuclear volume.

### Image analysis

#### Maximum cell thickness measurement

Confocal images were taken with 0.5μm steps in the z direction. Subsequently, the images were reconstructed using LAS X software (Leica microsystems, Wetzlar, Germany). Comparing x-z and z-y cross sections, the approximate region of interest (ROI) was found and subsequently the exact point that maximum thickness occurs were found enabling the thickness to be calculated.

#### Volume measurements

Captured images were analysed with Image J software. The area of each cell and nucleus has been measured and subsequently volume measured by integrating the cell area multiplied by the z stack thickness.

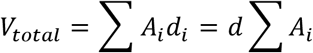

In which A*_i_* and d*_i_* are the area and thickness of the z stack number *i* respectively. The area was determined by setting a grey value for each cell and each z-stack image was carefully inspected and the area was approximated by free hand selection.

## Results and discussions

Many studies have shown that topographic cues on the substrate have a profound effect on cellular behaviour, cellular morphology, distribution and shape of internal cellular organelles (((29)(42),(43),(44)). Fabricated patterns on the micrometre scale provide a strong tool to study cell mechanics at the scale of forces that cells exert on themselves(45). Most of the studies in this field have focused on either the 2D morphological responses of the cells or mechanotransduction pathways. However it is only recently that 3D studies of the cell response to topographical cues has gained momentum. In this study, we try to give a clearer description of the effect of fibronectin stripes of different sizes on cellular and nuclear volume.

Fibronectin-printed stripes with different widths simulate oriented fibres in the underlying substrate. The oriented fibres occur in the form of oriented extracellular matrix (46)or aligned cytoskeletal fibres of the oriented cells underneath (47). Local and temporal polarization and orientation of the extracellular matrix, cytoskeleton and nucleus play a vital role in key cellular machinery such as cell division (48) and cell migration (20). In addition, cells both *in vivo* and *in vitro* frequently come into contact with polarized sections of adjacent or adhering cells below in the form of focal complexes (dot-like complexes of ~1×1 μm in size)(49), classical focal adhesions (streak like contacts of ~1×6 μm in size)(50), and supermature focal adhesions (highly elongated contacts of ~3×20 μm in size)(51). The interaction between cells at these contact points, which are joined by various molecular complexes made of CAMs play a key role in transducing mechanical signals to the cytoskeleton and nucleus(13). Due to the 3D nature of the interactions of cells in these regions, a comprehensive understanding of the transductions pathways will not be achieved without studying cells in the z-direction as well as in 2D.

## Maximum cell thickness

One of the features that plays a critical role in cellular function *in vivo* is cellular thickness. Cells don’t function as single layer of 2D sheets inside tissues; however, it is layer upon layer of the cells that form tissues and organs in the functioning body. As cells build up layers on top of each other, the dynamics of cellular thickness and how it depends on the substrate cues becomes important. *In vivo*, the substrate can consist of a layer of similar cells in the tissue, a layer of another formed by different cell type in adjacent tissue, a substrate formed by extracellular matrix fibre proteins or even a layer of medical synthetic prosthesis or synthetic tissue embedded inside the body of a patient. In all of these situations, understanding the dependency of cellular thickness dynamic on the underneath cue plays a vital role in understanding the whole tissue function and structure.

Fig. 2 shows the dependence of cell thickness on stripes of different width as well as a uniform control surface. This figure shows that the cell thickness in cells seeded on narrower fibronectin stripes (8 μm, 10μm and 12μm) is consistently higher than wider stripes (25μm, 50μm and evenly fibronectin coated substrate as the control)

**Fig. 2:**
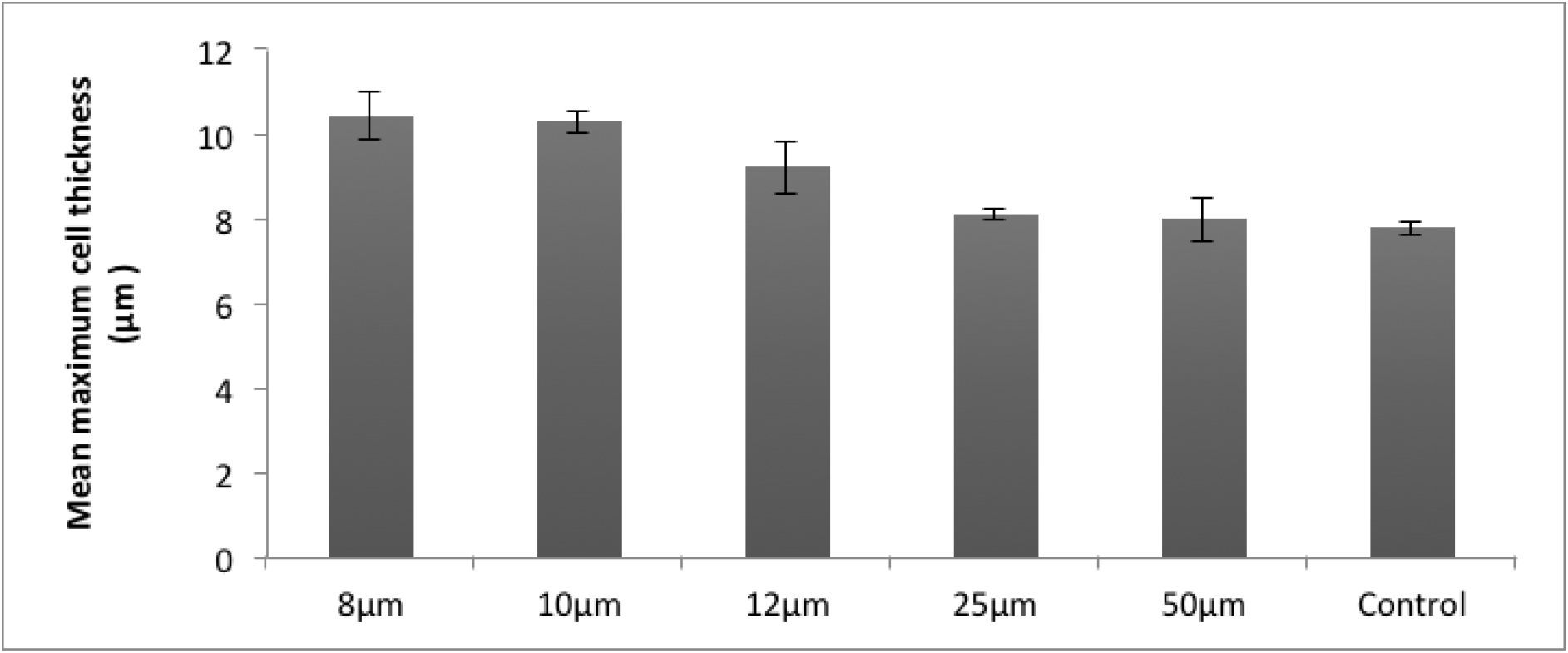
Maximum cell thickness among 3T3 cells spreading on 8μm (n=11), 10μm (n=79), 12 μm(n=15), 25μm(n=83) and 50μm(n=33) fibronectin stripes and evenly fibronectin coated substrate (n=87) on a PDMS substrate in the S-phase fixed with formaldehyde and stained with Phalloidin for F-actin. Error bars are SEM.

Fig. 3 shows stained cells in which the nucleus is shown in blue. On an unpatterned substrate the nucleus is not confined. On stripes of narrower and narrower width the ability of the nucleus to spread becomes increasingly constrained, so that on the 10mm stripe the nucleus becomes highly elongated. Thus a possible explanation for the difference in cell thickness seen in Fig. 2 could be the fact that stripes with width greater than the cell size (17.2μm ± 0.3μm s.e.m. (n=119)) or the nuclear width (13.5μm ± 0.2μm s.e.m. (n=142)) cannot effectively confine the nucleus and as a result, their effect on cellular thickness is minimal (Fig.3), whereas this is not the case for narrower stripes.

**Fig. 3:**
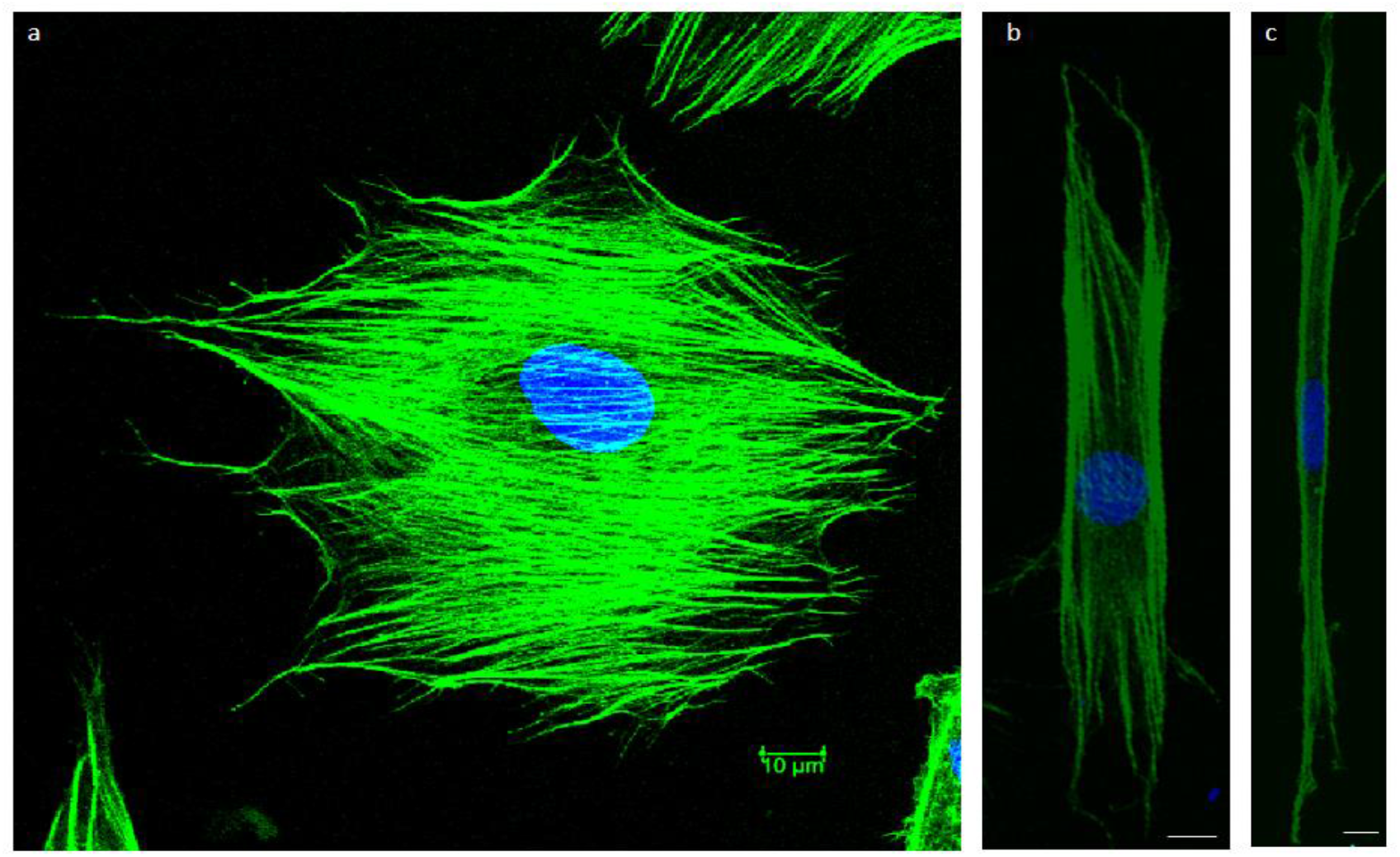
Nuclear confinement of the patterns. (**a**) Nucleus (blue) is not confined on a spreading cell on a uniformly fibronectin coated substrate. (**b**) Nucleus in still free to spread on cytoskeleton (green) and it is fairly symmetric on the fibroblast on a 25μm fibronectin coated stripe. (**c**) The nucleus is confined within highly elongated cytoskeleton and elongated itself in a cell spreading on a 10μm fibronectin coated stripe. Scale bar 10μm.

The aspect ratio (defined as major axis/minor axis ratio of the best fitting ellipse with the same area, orientation and centroid as the original cell/nucleus(52)) of the actin cytoskeleton and the nucleus also shows that although 25μm stripes can effectively impose contact guidance on actin cytoskeleton;, it doesn’t cause a significant elongation of the nucleus. On the other hand, narrower stripes such as those of 10μm width elongate both nucleus and actin cytoskeleton (Fig.4).

**Fig. 4:**
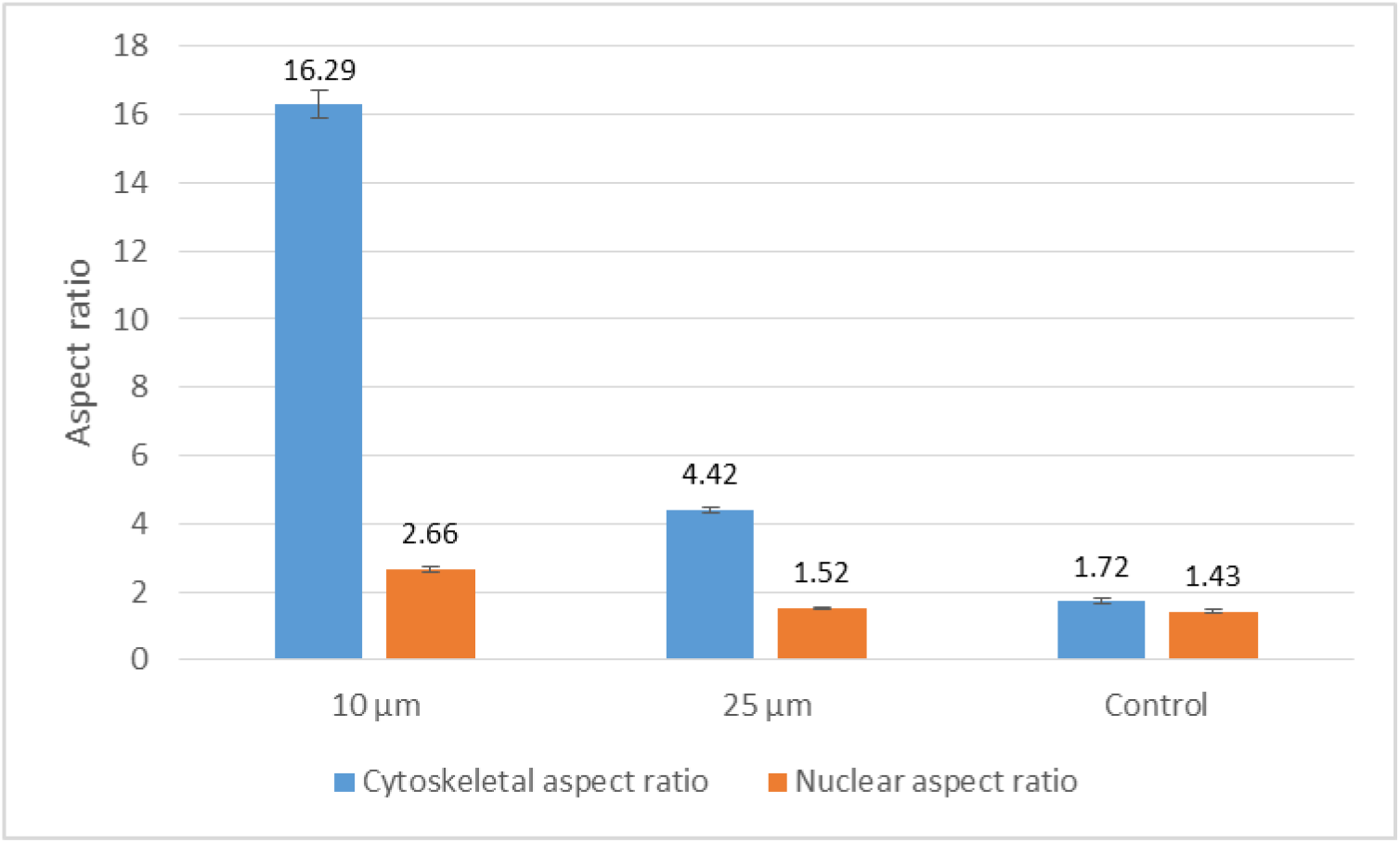
Aspect ratio of nucleus and actin cytoskeleton on 10μm (n=83, n_nucleus_=77), 25μm (n=79, n_nucleus_=82) and fibronectin stripes and evenly fibronectin coated control substrate (n=81, n_nucleus_=83) on a PDMS substrate in the S-phase fixed with formaldehyde and stained with Phalloidin for F-actin and TO-PRO-3 for nucleus. Error bars are SEM.

The forces that are exerted on the nucleus are either from conventional basal stress fibers or are from actin cap stress fibers((45),(53)). Conventional stress fibers are terminated by conventional focal adhesions (FA) complexes whereas actin cap stress fibers are terminated with actin cap associated fibers (ACAFs)(54). Actin cap stress fibers are composed of contractile bundles of actin filaments that are interconnected with phosphorylated myosin and α-actinin(20). These fibers are anchored to the top of the nucleus (Fig. 5(b)) and physically connected to the nucleus through the linker of nucleoskeleton and cytoskeleton (LINC) complexes (55) and they therefore play a crucial role in nuclear deformation and transducing mechanical signals to the nucleus (56). In fact among adherent cells, nuclear deformation is mainly governed by perinuclear actin contractile fibres rather than conventional basal stress fibers(57). It has been shown that for a significant proportion of the time, the actin cap is highly organized and elongated and strongly coupled to the nucleus; however for a small proportion of the time, the actin cap can be decoupled from the nucleus, allowing the nucleus to rotate and facilitate a migrating cell to re-orient itself in a new migratory direction(20). Both modelling and traction force experiments have shown that actin cap contractile fibers can exert forces of order of ~1−100 nN on the nucleus of adherent fibroblast cells ((58),(25),(59)). Considering the nuclear area of ~100 μm^2^ (nuclear area of detached cells=142 μm^2^±5 μm^2^ s.em. (n=142)), the perinuclear actin stress fibers can exert an effective pressure in order of ~ 1kP a which is enough to regulate the shape of a fibroblast nucleus with an approximate Young’s modulus of the order of ~1-10 kPa (60).

**Fig. 5:**
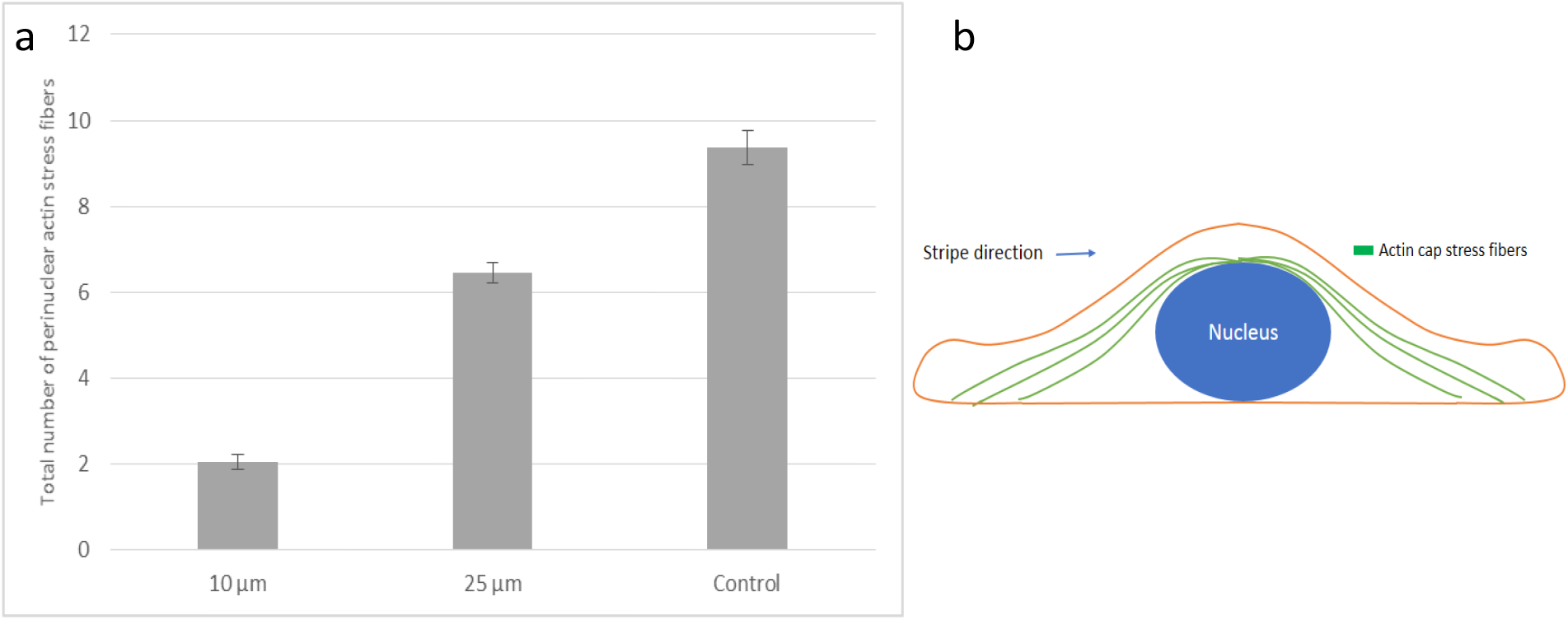
(**a**) The total number of perinuclear actin stress fibers among 3T3 cells spreading on 10μm (n=37), 25μm (n=30) and fibronectin stripes and evenly fibronectin coated control substrate (n=33) on a PDMS substrate in the S-phase fixed with formaldehyde and stained with Phalloidin for F-actin. Error bars are SEM. (**b**) Position of perinuclear actin cap stress fibers in a schematic cross-section of cell.

To explore the role of perinuclear actin stress fibers in determining cellular thickness adhering to different stripes, we counted the number of these fibers in each group of cells stained for filamentous actin.

Fig. 5 shows that cells seeded on narrower 10μm stripes have only about 20% of such contractile fibers compared to the cells spreading on control fibronectin coated substrate. The fibres can be seen in Fig. 6(a), while cells on wider 25μm stripes (Fig 6(b)) demonstrate almost about 60% of actin cap stress fibers were formed compared to cells adhering to the control fibronectin substrate. As Fig 6(c) shows, in unpolarised adhering cells, conventional basal stress fibers terminated by FA (yellow) at the cell area and are not well organized; however, actin cap associated stress fibers terminated by ACAFAs (red) at the periphery of the adherent cell and are highly organized. However in cells that are spreading on stripes, both conventional basal fibers and actin cap associated stress fibers are well organized and elongated in the direction of cell cytoskeleton (orange) and nucleus (blue) (Fig 6(d)). These results are another confirmation that there is a lower vertical compressive force acting on the nucleus of cells on narrower stripes to flatten the nucleus. (Fig.5).

**Fig. 6:**
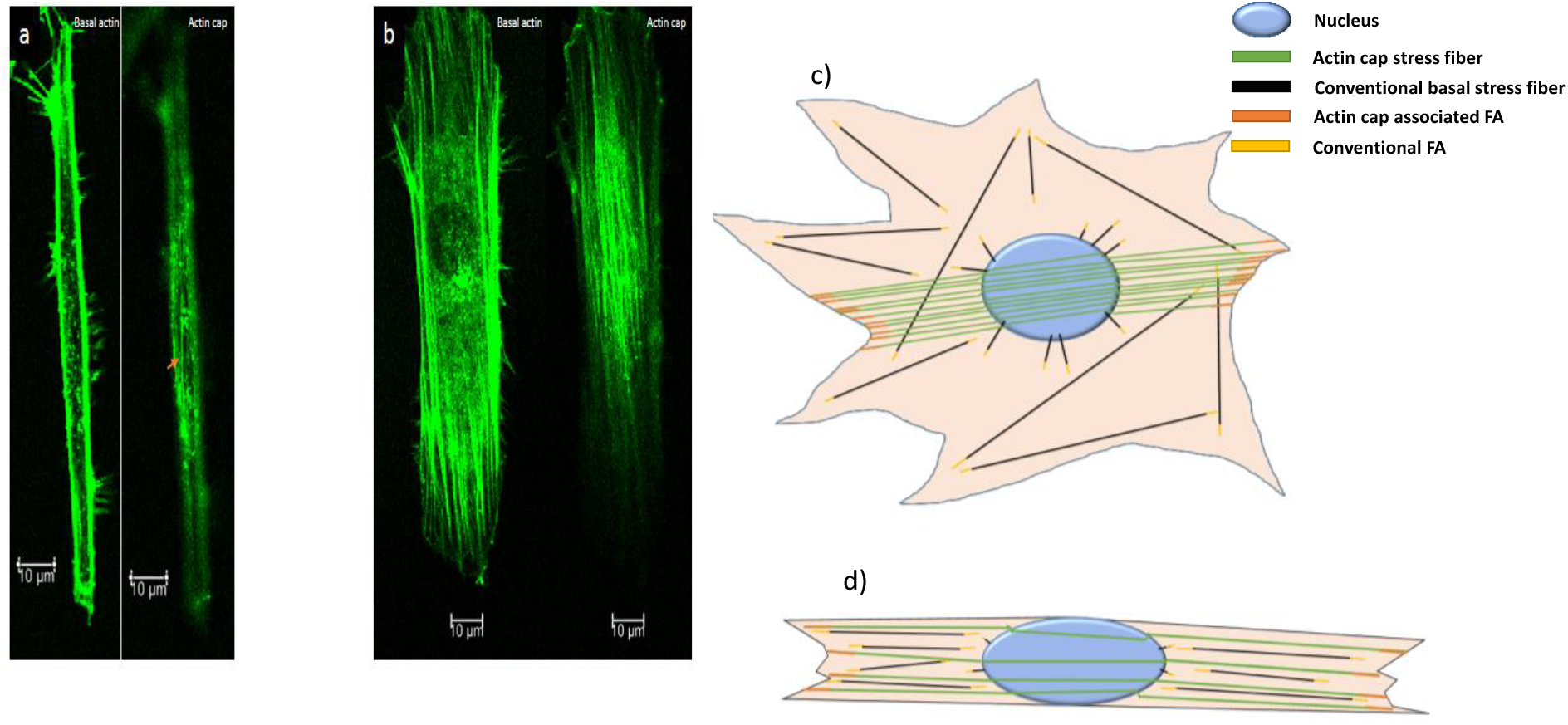
Actin cap stress fibers configuaration in adherent cells. (**a**)and (**b**) focal plane on basal surface reveals conventional basal stress fibers; wheras, focus on the apical plane on top of nucleus reveals highly organized perinuclar actin cap stress fibers among 3T3 fibroblast cells on 8μm (**a**) and 25μm(**b**) fibronectin stripes. The only actin cap stress fiber in the cell adherent to 8μm pattern cell is marked with a red arrow. (**c**) Schematic of stress fibers configuration in an unpolarised adhering cell. This schematic image was inspired by (61)(**d**) Schematic of stress fibers configuration in a polarized adherent cell.

### Cellular volume

The volume of cells is a critical parameter in the determination of organ and tissue morphology and thereby in the main function of living organisms. A cell’s ability to regulate its volume plays a central role in cell function. Volume regulation gives the cell protective and adaptive advantages by allowing cytoskeletal rearrangements(62). In addition, changes in cellular volume triggers signalling pathways for cell proliferation, death, and migration (63). Vertebrate cells, with a few exceptions, are permeable to water and lack the stiff cellular wall of plants and bacteria(62). The cell volume is controlled by the interplay of membrane tension, active contractility and water/ion influx(63). It has been shown previously that adherent mouse embryonic fibroblasts have a bigger volume compared to detached cells (25).

Fig. 7 shows that cell volume almost doubles during the adherence process from an initially detached state. However as the cell is being confined within fibronectin stripes, the volume increase is partly inhibited. As the pattern gets wider, the cellular volume becomes comparable to that of cells adhering to unpatterned fibronectin surfaces. It is well known that actin polymerizing is the main force driving morphological changes during cell spreading in the adhesion process(64).We hypothesize that confining cells to the patterns inhibits this actin polymerization process in the perpendicular direction to the stripe’s direction and this way, reduces the total cellular volume.

**Fig. 7:**
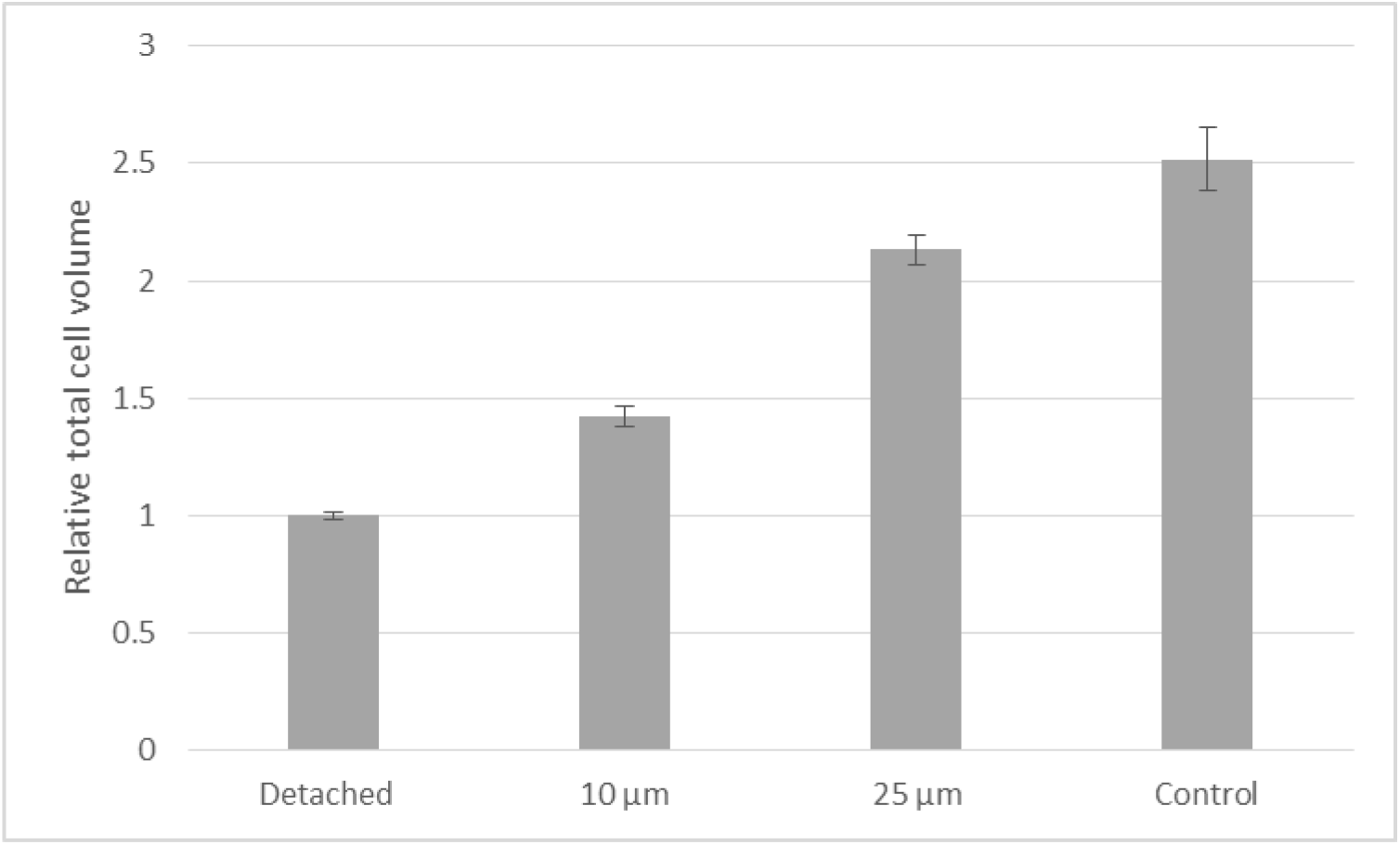
Relative total cell volume change on fibronectin stripes on 10μm (n=81), 25μm (n=79) and fibronectin stripes and evenly fibronectin coated control substrate (n=82) on a PDMS substrate in the S-phase fixed with formaldehyde and stained with Phalloidin for F-actin and suspended detached cells estimated by bright field microscopy (n=119). Error bars are SEM.

### Nuclear deformation

The nucleus is the stiffest and largest organelle in eukaryotic cells and contains most of the genetic information of the cell(65). It is the site for the major functions of a cell such as DNA replication, transcriptional regulations, and RNA processing and ribosome maturation(25). The lamina envelope of nucleus is coupled with the cytoskeleton via a series of binding proteins to the actin and intermediate filaments (66).Therefore, the intracellular and extracellular forces affect the shape and structure of the nucleus and inevitably the cell signalling and gene transcription. It has been shown that cytoskeletal motors actively control the nucleus shape. Microtubule motors apply a compressive pressure, whereas actin stress fibers apply compressive force on the nucleus (25). The substrate topography and stiffness can deform the nucleus (45),(29)).

Fig. 8 shows that the nuclear volume almost doubles during cell adherence in comparison to the nucleus of detached cells. Kim et al (25) carried out an experimental and modelling study of the dynamics of nuclear shape during detachment process which is consistent with our results. They concluded that the volume shrinkage in the detachment process is a result of three factors; the highly folded nucleus in detached cells, pressure difference across the nuclear envelope and cytoplasmic mechanical forces.

**Fig. 8:**
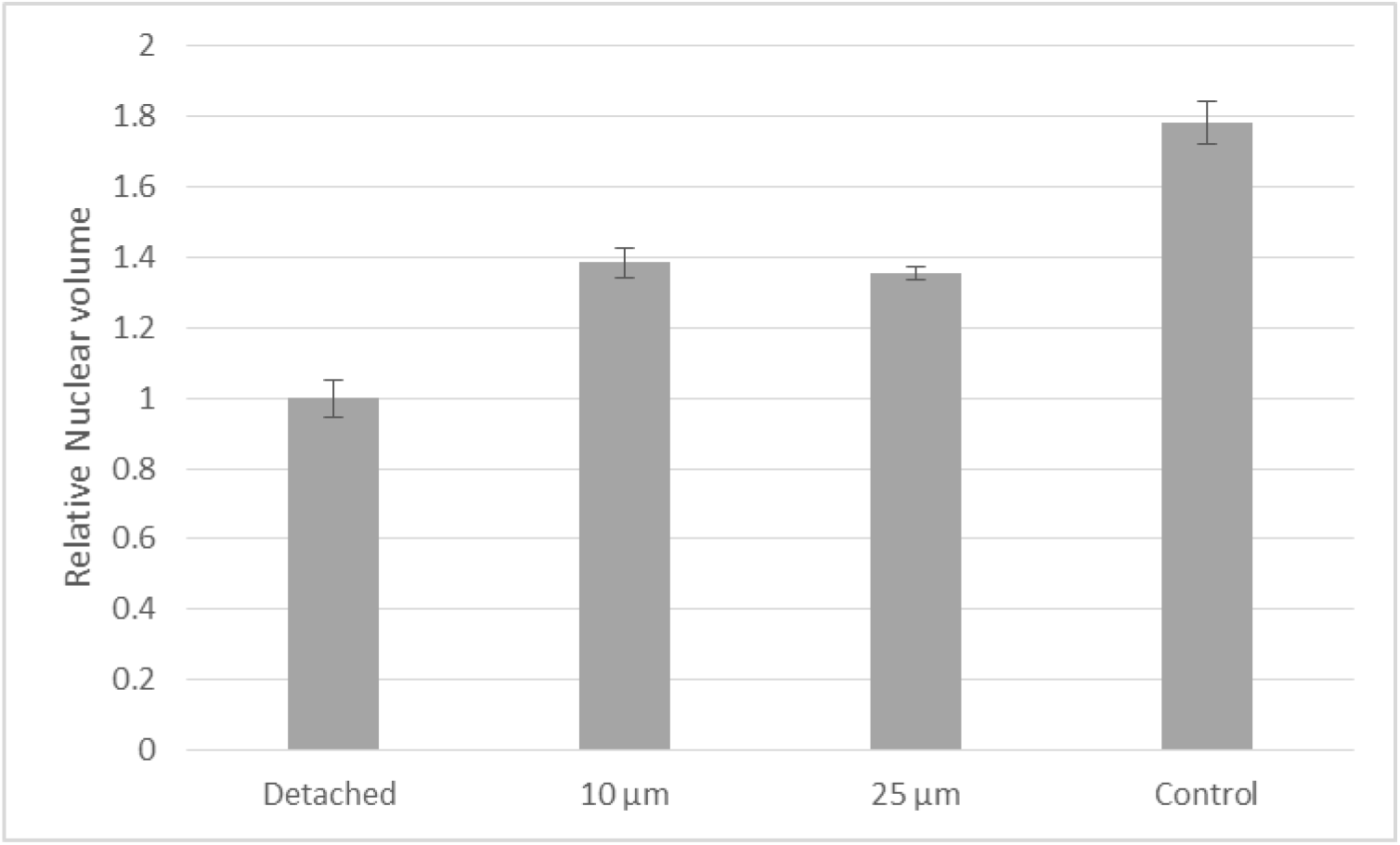
Relative nuclear volume change on fibronectin stripes on 10μm (n=82), 25μm (n=84) and fibronectin stripes and evenly fibronectin coated control substrate (n=91) on a PDMS substrate in the S-phase fixed with formaldehyde and stained with Phalloidin for F-actin and suspended detached cells stained with Hoechst 33342 (n=142). Error bars are SEM.

Our results further shows that confining cells to fibronectin stripes limits this nuclear volume increase. However, the size of fibronectin stripes did not affect the nuclear volume considerably despite the clear difference in nuclear aspect ratio between 10μm and 25μm (Fig.4). This might result from the fact that the force from basal contractile fibers has more effect on nucleus volume and these fibers might act similarly on pattern with different sizes(45).

We also suspect that differences in the distribution of the cell cycle for different patterns can play a role in this result as we came across more dividing cells adhering to the control substrate. Previous studies have shown that cell volume change during cell cycle and almost doubles during interphase of dividing cells (67). Unfortunately, it was not always easy to distinguish a dividing nucleus from a slightly elongated one. Furthermore, our staining was not sufficient to correlate nuclear volume with cell cycle. We think that further studies is needed to understand how the fibronectin stripes might affect cell cycle.

## Conclusion

In this study we measured the morphological responses of NIH/3T3 mouse embryonic fibroblast’s cytoskeleton and nucleus to fibronectin stripes printed with different widths on a PDMS substrate. We observed that stripe size significantly affects cytoskeletal and nuclear morphology. In particular we found that stripes narrower than the size scale of unattached fibroblast before adhesion fibroblasts have the most effect on the cellular morphology and confinement to the stripes. Our results indicate that cells adhering to narrower stripes are thicker and have got a more elongated cytoskeleton and nucleus in comparison to the cells which were seeded on unpatterned fibronection coated substrates. In addition, these cells have got fewer actin contractile fibers on the top of their nucleus. Furthermore, we have shown that the total cellular volume increases during cell adhesion processe and there is a correlation between volume increase and the width of stripes. During cell adhesion, the nuclear volume also increased significantly; however, confinement to the fibronectin stripes reduced nuclear volume expansion independent from the of stripe size. Our results give a quantified insight into the interaction of cells with a topography that mimics frequently occurring situations in the living microenvironment of the adherent cells. Our results helps to gain better understanding of the mechanobiolgy of the cell. In addition, our quantified approach can be a valuable tool to predict and potentially control the cell-substrate interaction which is highly applicable in the field of tissue engineering and regenerative medicine.

## Acknowledgments

Authors thank Dr. Otti Croze, Professor Piettro. Cicuta, Dr. Lorenzo Michele and Dr. Christopher Lester for helpful advices and useful discussions about the topic. Pooya Mamaghani thanks Fiona Morgan and Dr. Jonny Huang for providing lab trainings and supports. He also thanks the Cambridge Overseas Trust, the Islamic Development Bank, Cavendish Laboratory and Peterhouse College for funding his PhD course.

## References

1. Gumbiner BM. Cell adhesion: the molecular basis of tissue architecture and morphogenesis. Cell. 1996 Mar 9;84(3):345–57.

2. Weiss P. In vitro experiments on the factors determining the course of the outgrowing nerve fiber. J Exp Zool. 1934 Aug;68(3):393–448.

3. Alberts BJALJRM coaut. coaut. coaut. BAAJJLMRKRPW. Molecular biology of the cell. 2008.

4. Cassimeris L, Lingappa VR, Plopper G, Lewin B. Lewin’s cells. Jones and Bartlett Publishers; 2011. 1053 p.

5. Marmaras A, Lendenmann T, Civenni G, Franco D, Poulikakos D, Kurtcuoglu V, et al. Topography-mediated apical guidance in epidermal wound healing. Soft Matter. 2012;8(26):6922.

6. Stuart ES, Moscona AA. Embryonic morphogenesis: role of fibrous lattice in the development of feathers and feather patterns. Science. 1967 Aug 25;157(3791):947–8.

7. Hoffman-Kim D, Mitchel JA, Bellamkonda R V. Topography, cell response, and nerve regeneration. Annu Rev Biomed Eng. 2010 Aug 15;12:203–31.

8. Bauer AL, Jackson TL, Jiang Y. Topography of Extracellular Matrix Mediates Vascular Morphogenesis and Migration Speeds in Angiogenesis. Czirók A, editor. PLoS Comput Biol. 2009 Jul 24;5(7):e1000445.

9. Song S, Kim EJ, Bahney CS, Miclau T, Marcucio R, Roy S. The synergistic effect of micro-topography and biochemical culture environment to promote angiogenesis and osteogenic differentiation of human mesenchymal stem cells. Acta Biomater. 2015;18:100–11.

10. Yim EKF, Darling EM, Kulangara K, Guilak F, Leong KW. Nanotopography-induced changes in focal adhesions, cytoskeletal organization, and mechanical properties of human mesenchymal stem cells. Biomaterials. 2010 Feb;31(6):1299–306.

11. Tilghman RW, Parsons JT. Focal adhesion kinase as a regulator of cell tension in the progression of cancer. Semin Cancer Biol. 2008 Feb;18(1):45–52.

12. Geiger B, Spatz JP, Bershadsky AD. Environmental sensing through focal adhesions. Nat Rev Mol Cell Biol. 2009 Jan;10(1):21–33.

13. Vogel V, Sheetz M. Local force and geometry sensing regulate cell functions. Nat Rev Mol Cell Biol. 2006 Apr 22;7(4):265–75.

14. Engler AJ, Sen S, Sweeney HL, Discher DE. Matrix elasticity directs stem cell lineage specification. Cell. 2006 Aug 25;126(4):677–89.

15. Delmas P, Hao J, Rodat-Despoix L. Molecular mechanisms of mechanotransduction in mammalian sensory neurons. Nat Rev Neurosci. 2011 Mar 9;12(3):139–53.

16. Ohara PT, Buck RC. Contact guidance in vitro. Exp Cell Res. 1979 Jul 1;121(2):235–49.

17. Nikkhah M, Edalat F, Manoucheri S, Khademhosseini A. Engineering microscale topographies to control the cell–substrate interface. Biomaterials. 2012 Jul;33(21):5230–46.

18. Stevenson PM, Donald AM. Identification of three regimes of behavior for cell attachment on topographically patterned substrates. Langmuir. 2009 Jan 6;25(1):367–76.

19. Huang C-K, Donald A. Revealing the dependence of cell spreading kinetics on its spreading morphology using microcontact printed fibronectin patterns. J R Soc Interface. 2014 Dec 3;12(102):20141064–20141064.

20. Kim D-H, Cho S, Wirtz D. Tight coupling between nucleus and cell migration through the perinuclear actin cap. J Cell Sci. 2014 Jun 1;127(11):2528–41.

21. Oakley C, Brunette DM. The sequence of alignment of microtubules, focal contacts and actin filaments in fibroblasts spreading on smooth and grooved titanium substrata. J Cell Sci. 1993 Sep;106 (Pt 1:343–54.

22. Kung KS, Canton I, Massignani M, Battaglia G, Donald AM. The development of anisotropic behaviours of 3T3 fibroblasts on microgrooved patterns. Eur Phys J E Soft Matter. 2011 Mar;34(3):23.

23. Kasten A, Naser T, Brüllhoff K, Fiedler J, Müller P, Möller M, et al. Guidance of Mesenchymal Stem Cells on Fibronectin Structured Hydrogel Films. Engler AJ, editor. PLoS One. 2014 Oct 15;9(10):e109411.

24. Khatau SB, Hale CM, Stewart-Hutchinson PJ, Patel MS, Stewart CL, Searson PC, et al. A perinuclear actin cap regulates nuclear shape. Proc Natl Acad Sci. 2009 Nov 10;106(45):19017–22.

25. Kim D-H, Li B, Si F, Phillip JM, Wirtz D, Sun SX. Volume regulation and shape bifurcation in the cell nucleus. J Cell Sci. 2015 Sep 15;128(18):3375–85.

26. Chalut KJ, Kulangara K, Giacomelli MG, Wax A, Leong KW. Deformation of stem cell nuclei by nanotopographical cues. Soft Matter. 2010 Apr 21;6(8):1675–81.

27. Hanson L, Zhao W, Lou H-Y, Lin ZC, Lee SW, Chowdary P, et al. Vertical nanopillars for in situ probing of nuclear mechanics in adherent cells. Nat Nanotechnol. 2015 Jun;10(6):554–62.

28. Pan Z, Yan C, Peng R, Zhao Y, He Y, Ding J. Control of cell nucleus shapes via micropillar patterns. Biomaterials. 2012 Feb;33(6):1730–5.

29. Badique F, Stamov DR, Davidson PM, Veuillet M, Reiter G, Freund J-N, et al. Directing nuclear deformation on micropillared surfaces by substrate geometry and cytoskeleton organization. Biomaterials. 2013 Apr;34(12):2991–3001.

30. Chalut KJ, Kulangara K, Giacomelli MG, Wax A, Leong KW. Deformation of stem cell nuclei by nanotopographical cues. Soft Matter. 2010 Apr;6(8):1675–81.

31. Kulangara K, Adler AF, Wang H, Chellappan M, Hammett E, Yasuda R, et al. The effect of substrate topography on direct reprogramming of fibroblasts to induced neurons. Biomaterials. 2014 Jul;35(20):5327–36.

32. Kim D-H, Cho S, Wirtz D. Tight coupling between nucleus and cell migration through the perinuclear actin cap. J Cell Sci. 2014 Jun;127(11):2528–41.

33. Qin D, Xia Y, Whitesides GM. Soft lithography for micro- and nanoscale patterning. Nat Protoc. 2010 Mar;5(3):491–502.

34. Shen K, Qi J, Kam LC. Microcontact printing of proteins for cell biology. J Vis Exp. 2008 Dec 5;(22).

35. Tan SH, Nguyen N-T, Chua YC, Kang TG. Oxygen plasma treatment for reducing hydrophobicity of a sealed polydimethylsiloxane microchannel. Biomicrofluidics. 2010 Sep 30;4(3):32204.

36. Döbereiner H-G, Dubin-Thaler B, Giannone G, Xenias HS, Sheetz MP. Dynamic Phase Transitions in Cell Spreading. Phys Rev Lett. 2004 Sep 2;93(10):108105.

37. Cuvelier D, Théry M, Chu Y-S, Dufour S, Thiéry J-P, Bornens M, et al. The Universal Dynamics of Cell Spreading. Curr Biol. 2007 Apr;17(8):694–9.

38. Mooney DJ, Langer R, Ingber DE. Cytoskeletal filament assembly and the control of cell spreading and function by extracellular matrix. J Cell Sci. 1995 Jun;108 (Pt 6):2311–20.

39. Fardin MA, Rossier OM, Rangamani P, Avigan PD, Gauthier NC, Vonnegut W, et al. Cell spreading as a hydrodynamic process. Soft Matter. 2010 Aug 10;6:4788–99.

40. Xiong Y, Rangamani P, Fardin M-A, Lipshtat A, Dubin-Thaler B, Rossier O, et al. Mechanisms Controlling Cell Size and Shape during Isotropic Cell Spreading. Biophys J. 2010 May 19;98(10):2136–46.

41. Fujiwara I, Takahashi S, Tadakuma H, Funatsu T, Ishiwata S. Microscopic analysis of polymerization dynamics with individual actin filaments. Nat Cell Biol. 2002 Sep 19;4(9):666–73.

42. Anselme K, Bigerelle M. Role of materials surface topography on mammalian cell response. Int Mater Rev. 2011 Jul 12;56(4):243–66.

43. S. Yamamoto *,†,‡, M. Tanaka ‡,§, H. Sunami †,‡, E. Ito ‡, S. Yamashita ‡, Y. Morita † and, et al. Effect of Honeycomb-Patterned Surface Topography on the Adhesion and Signal Transduction of Porcine Aortic Endothelial Cells. 2007;

44. Anselme K, Ponche A, Bigerelle M. Relative influence of surface topography and surface chemistry on cell response to bone implant materials. Part 2: Biological aspects. Tanner KE, Dalby MJ, editors. Proc Inst Mech Eng Part H J Eng Med. 2010 Dec 29;224(12):1487–507.

45. Hanson L, Zhao W, Lou H-Y, Lin ZC, Lee SW, Chowdary P, et al. Vertical nanopillars for in situ probing of nuclear mechanics in adherent cells. Nat Nanotechnol. 2015 Jun 18;10(6):554–62.

46. Jia S, Liu L, Pan W, Meng G, Duan C, Zhang L, et al. Oriented cartilage extracellular matrix-derived scaffold for cartilage tissue engineering. J Biosci Bioeng. 2012 May;113(5):647–53.

47. Zolessi FR, Poggi L, Wilkinson CJ, Chien C-B, Harris WA. Polarization and orientation of retinal ganglion cells in vivo. Neural Dev. 2006 Oct;1(1):2.

48. Théry M, Racine V, Pépin A, Piel M, Chen Y, Sibarita J-B, et al. The extracellular matrix guides the orientation of the cell division axis. Nat Cell Biol. 2005 Oct 18;7(10):947–53.

49. Beningo KA, Dembo M, Kaverina I, Small JV, Wang Y. Nascent Focal Adhesions Are Responsible for the Generation of Strong Propulsive Forces in Migrating Fibroblasts. J Cell Biol. 2001;153(4).

50. Goffin JM, Pittet P, Csucs G, Lussi JW, Meister J-J, Hinz B. Focal adhesion size controls tension-dependent recruitment of alpha-smooth muscle actin to stress fibers. J Cell Biol. 2006 Jan 16;172(2):259–68.

51. Goffin J, Csucs G, Lussi J, Meister J, Hinz B. Masters and Servants of the Force: Focal Adhesion Size Controls Recruitment of α-Smooth Muscle Actin to Stress Fibers. Wound Repair Regen. 2005 Jan 17;13(1):A20–A20.

52. Weblet Importer [Internet]. Available from: https://imagej.nih.gov/ij/source/ij/process/EllipseFitter.java

53. Khatau SB, Hale CM, Stewart-Hutchinson PJ, Patel MS, Stewart CL, Searson PC, et al. A perinuclear actin cap regulates nuclear shape. Proc Natl Acad Sci. 2009 Nov;106(45):19017–22.

54. Kim D-H, Khatau SB, Feng Y, Walcott S, Sun SX, Longmore GD, et al. Actin cap associated focal adhesions and their distinct role in cellular mechanosensing. Sci Rep. 2012 Dec 3;2(1):555.

55. Rothballer A, Schwartz TU, Kutay U. LINCing complex functions at the nuclear envelope. Nucleus. 2013 Jan 28;4(1):29–36.

56. Kim D-H, Chambliss AB, Wirtz D. The multi-faceted role of the actin cap in cellular mechanosensation and mechanotransduction. Soft Matter. 2013 Jun 21;9(23):5516–23.

57. Tamiello C, Bouten CVC, Baaijens FPT. Competition between cap and basal actin fiber orientation in cells subjected to contact guidance and cyclic strain. Sci Rep. 2015 Mar 4;5:8752.

58. Munevar S, Wang Y, Dembo M. Traction force microscopy of migrating normal and H-ras transformed 3T3 fibroblasts. Biophys J. 2001 Apr;80(4):1744–57.

59. Sabass B, Gardel ML, Waterman CM, Schwarz US. High resolution traction force microscopy based on experimental and computational advances. Biophys J. 2008 Jan 1;94(1):207–20.

60. Ferrera D, Canale C, Marotta R, Mazzaro N, Gritti M, Mazzanti M, et al. Lamin B1 overexpression increases nuclear rigidity in autosomal dominant leukodystrophy fibroblasts. FASEB J. 2014 Sep 1;28(9):3906–18.

61. Kim D-H, Chambliss AB, Wirtz D. The multi-faceted role of the actin cap in cellular mechanosensation and mechanotransduction. Soft Matter. 2013 Jun 21;9(23):5516–23.

62. Hoffmann EK, Lambert IH, Pedersen SF. Physiology of cell volume regulation in vertebrates. Physiol Rev. 2009 Jan 1;89(1):193–277.

63. Jiang H, Sun SX. Cellular pressure and volume regulation and implications for cell mechanics. Biophys J. 2013 Aug 6;105(3):609–19.

64. Parsons JT, Horwitz AR, Schwartz MA. Cell adhesion: integrating cytoskeletal dynamics and cellular tension. Nat Rev Mol Cell Biol. 2010 Sep;11(9):633–43.

65. Dahl KN, Ribeiro AJS, Lammerding J. Nuclear shape, mechanics, and mechanotransduction. Circ Res. 2008 Jun 6;102(11):1307–18.

66. Herrmann H, Bär H, Kreplak L, Strelkov S V, Aebi U. Intermediate filaments: from cell architecture to nanomechanics. Nat Rev Mol Cell Biol. 2007 Jul;8(7):562–73.

67. Maeshima K, Iino H, Hihara S, Imamoto N. Nuclear size, nuclear pore number and cell cycle. Nucleus. 2011 Mar 28;2(2):113–8.

